# *Escherichia coli* outer membrane vesicles encapsulating small molecule antibiotics improve drug function by facilitating transport

**DOI:** 10.1101/2025.01.08.631972

**Authors:** Meishan Wu, Rachael M. Harrower, Ziang Li, Angela C. Brown

## Abstract

The development of novel antimicrobial agents that are effective against Gram-negative bacteria is hindered by the dual membrane cell wall structure of these bacteria. To reach their intracellular targets, most antibiotics must therefore pass through protein channels called porins; however, a common mechanism of acquired resistance is decreased expression of these proteins. Inspired by the ability of outer membrane vesicles (OMVs) to deliver cargo to the bacterial cytosol, we hypothesized that encapsulation of small molecule antibiotics within OMVs would improve the activity of the drugs by facilitating uptake. To test this, we investigated the ability of antibiotic- encapsulated OMVs to inhibit the growth of both Gram-negative and Gram-positive bacteria, including multidrug-resistant clinical isolates. We also demonstrated that this mechanism of delivery does not require porin expression. Together, our results demonstrate the potential of OMVs as novel antibiotic delivery vehicles to treat antibiotic-resistant bacterial infections.

---

The emergence and rapid spread of antibiotic resistance presents an alarming threat to global health. Among the list of 18 antibiotic-resistant bacteria and fungi identified by the United States Centers for Disease Control and Prevention, 11 are Gram-negative bacteria^1,2^. These bacteria exhibit resistance to a wide range of antibiotics, and effective treatment options are scarce. The last class of broad-spectrum antibiotics to be developed was the quinolone family, which was first introduced in the 1960s^3^. This lack of new drugs has led to a critical shortage of viable treatment options for infections caused by these resistant strains, with clinicians relying on older drugs such as colistin, that are associated with severe adverse effects^4^.

The primary factor leading to limited activity of many antibiotics against Gram-negative bacteria is their impenetrable cell envelope, which is composed of an outer membrane and an inner membrane, separated by a periplasmic space. The outer membrane is rich in lipopolysaccharide (LPS), which serves as a protective barrier against harmful substances, such as antibiotics. Beneath the outer membrane lies the periplasmic space, filled with a gel-like matrix containing a peptidoglycan layer that provides structural support to the cell. The inner membrane is composed of a phospholipid bilayer that regulates the transport of ions and molecules, further limiting uptake of antibiotics^5,6^. In order to reach their cytosolic targets, antibiotics must therefore pass through protein channels called porins. However, large antibiotics, such as glycopeptides are unable to pass through these porins^7,8^, and bacteria can downregulate porin expression to limit drug uptake^9–13^.

Recent studies have therefore focused on enhancing antibiotic effectiveness by combining antibiotics with certain adjuvants to promote rapid drug uptake using membrane permeabilizers or lipid-based nanodelivery platforms. Membrane permeabilizers, such as polymyxin B and ethylenediaminetetraacetic acid (EDTA), work by disrupting the structure of the bacterial outer membrane^14,15^. When used in combination with antibiotics, membrane permeabilizers improve the transport of antibiotics across the outer membrane by increasing the fluidity of the lipid bilayer^16,17^. However, the use of permeabilizers *in vivo* is limited by their adverse effects on mammalian cells, as the molecules can also affect the host cell membranes^18,19^. An alternative approach to improve antibiotic delivery involves the use of lipid-based delivery systems, such as liposomes. These carriers are designed with specific lipid compositions that enable them to fuse with the bacterial outer membrane^20,21^. Such fusogenic liposomes have been successfully employed to broaden the spectrum of antibiotics, such as vancomycin, which is otherwise ineffective against Gram-negative bacteria^22^. Despite their promise, these lipid-based nanodelivery systems face challenges related to their physical instability, which can lead to drug leakage over time^23,24^.

Gram-negative bacteria release vesicles derived from their outer membrane, called “outer membrane vesicles” (OMVs) throughout all stages of growth^25,26^. OMVs are capable of encapsulating and delivering nucleic acids, signaling molecules, enzymes, and toxins to promote the survival of parent bacteria^27–31^. Importantly, these vesicles enable content delivery to bacterial cells by fusing with their outer membrane^32^ allowing the transfer of genetic material or other macromolecules directly into the recipient cells^33–37^. In addition, the stability of OMVs ensures the protection of their cargo in harsh environments^38,39^.

We therefore proposed that OMVs might promote antibiotic delivery across the Gram-negative cell membrane, thereby enhancing the effectiveness of these drugs. We have previously explored several methods for the loading of fluoroquinolones into the lumen of *Escherichia coli* OMVs. Our results demonstrated that (1) small molecules can be readily loaded into *E. coli* OMVs, and (2) active loading methods, such as electroporation and sonication, can efficiently load antibiotics into OMVs with an encapsulation efficiency reaching as high as 68%^40^.

In this study, we explored the antibacterial potential of antibiotic-loaded OMVs (aOMVs) against laboratory and multidrug-resistant (MDR) clinical isolates of Gram-negative and Gram-positive bacteria by comparing the activity of aOMVs, free (unencapsulated) antibiotics, and empty OMVs. To demonstrate that OMVs enable delivery via a porin-independent pathway, we tested aOMVs on porin-deficient single-gene knockout mutants of *E. coli*. Together, our results demonstrate that by improving delivery across the Gram-negative outer membrane, in a porin-independent manner, OMVs are able to enhance the activity of small molecule antibiotics against a range of bacterial species.

## Results

Characterization and stability of aOMVs The morphology of the empty OMVs, determined using scanning electron microscopy (SEM), is displayed in Fig. 1. The empty OMVs appeared spherical, with an average diameter of 33.7 ± 7.0 nm (n=25). The sizes of the OMVs and aOMVs measured using dynamic light scattering (DLS) are reported in Table 1. The measured diameter of the untreated OMVs is consistent with those observed using SEM. No significant difference in size was observed between the unloaded OMVs and those loaded with imipenem (IMI).

**Figure 1:**
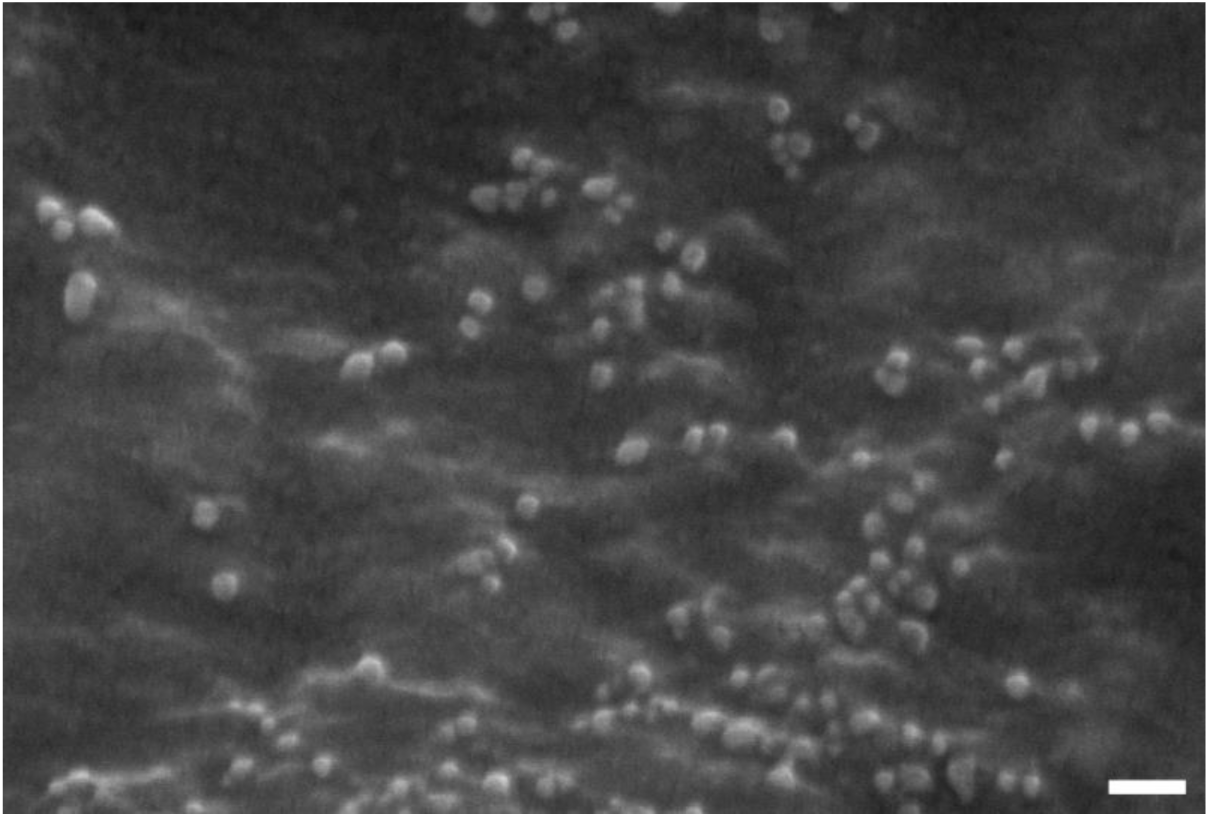
***E. coli* JC8031 OMVs imaged using SEM.** Unloaded OMVs are spherical and homogeneous in size with an average diameter of 33.7 ± 7.0 nm (n=25) analyzed using ImageJ. Scale bar = 100 nm.

**Table 1:** Average diameter of OMVs and aOMVs measured using DLS.

| *Condition* |  | *Diameter (nm)* |
| --- | --- | --- |
| Untreated | Empty OMVs | $38.33 \pm 3.64$ |
| | IMI-OMVs | $40.38 \pm 3.29$ |
| Post-incubation<br>(24 hr, 37 °C) | Empty OMVs | $42.98 \pm 2.33$ |
| | IMI-OMVs | $39.64 \pm 1.46$ |

The diameter represents the mean (n=3) ± standard deviation.

Sonication was used to synthesize aOMVs, and the encapsulation efficiencies (EEs) for IMI-loaded OMVs (IMI-OMVs) was found to be 41% ± 2.39%. We also investigated OMV stability by incubating the aOMVs at 37 °C for 24 hr. We showed, using DLS, that the size of the OMVs did not change upon incubation for 24 hr (Table 1); in addition, the content leakage from IMI-OMVs was determined to be less than 1% of the total amount encapsulated. These results indicate that OMVs are resistant to thermal degradation and can provide a stable environment for cargo.

The diameter represents the mean (n=3) ± standard deviation.

### 1. 2. Delivery of aOMVs to Gram-negative bacteria

We previously showed that encapsulation of IMI in OMVs enhances the antibacterial activity of this drug against a lab strain of *E. coli*, W3110^40^. To demonstrate the wider applicability of the approach, we tested the effectiveness of IMI-OMVs against a lab strain of *Pseudomonas aeruginosa,* PAO1. PAO1 cells were treated with free IMI, IMI-OMVs (with the same IMI concentration) or empty OMVs (with the same OMV concentration). Bacterial growth was determined by following the optical density (OD600) over time. The half-maximal normalized absorbance of the PBS-treated control (IC50), served as a reference, where values below the IC50 were determined to represent “effective” concentrations. We observed that IMI-OMVs were as effective as free IMI in inhibiting the growth of this strain of bacteria (Fig. 2, Fig. S1), with both IMI and IMI-OMVs crossing the IC50 at a concentration of 1 µg/mL.

**Figure 2:**
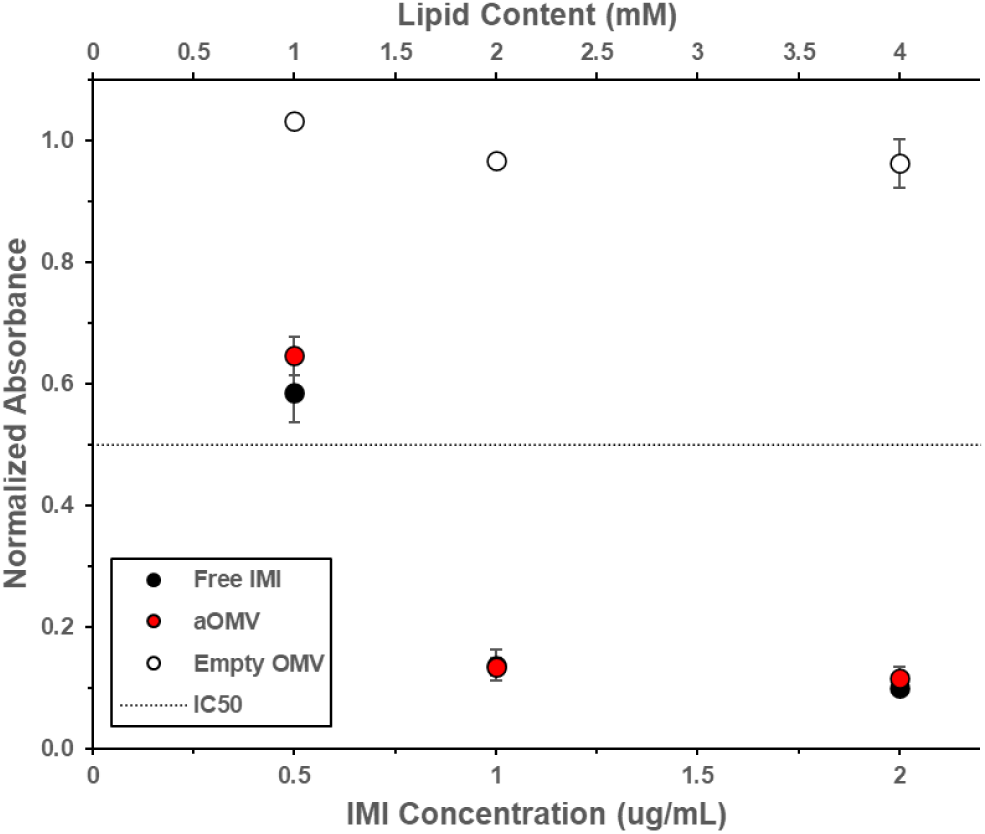
Delivery of IMI, both free and encapsulated within OMVs, to *P. aeruginosa* strain PAO1. Bacteria were treated with free IMI (black), IMI-OMVs (red), or empty OMVs (white). The change in absorbance at 600 nm after 12 hr of growth of treated bacteria was normalized to that of untreated bacteria. Each data point represents the mean (n=3) ± standard deviation. The dashed line represents the IC50.

Next, we furthered these studies by investigating the activity of aOMVs in MDR clinical isolates of *P. aeruginosa*, obtained from the CDC/FDA Antimicrobial Resistance Isolate Bank. We measured the susceptibility of three selected isolates to unloaded IMI, IMI-OMVs, and empty OMVs. Fig. 3 shows the change in absorbance after 12 hr for bacteria treated with either free IMI, IMI-OMVs, or empty OMVs, normalized by the change in absorbance of untreated bacteria.

**Figure 3:**
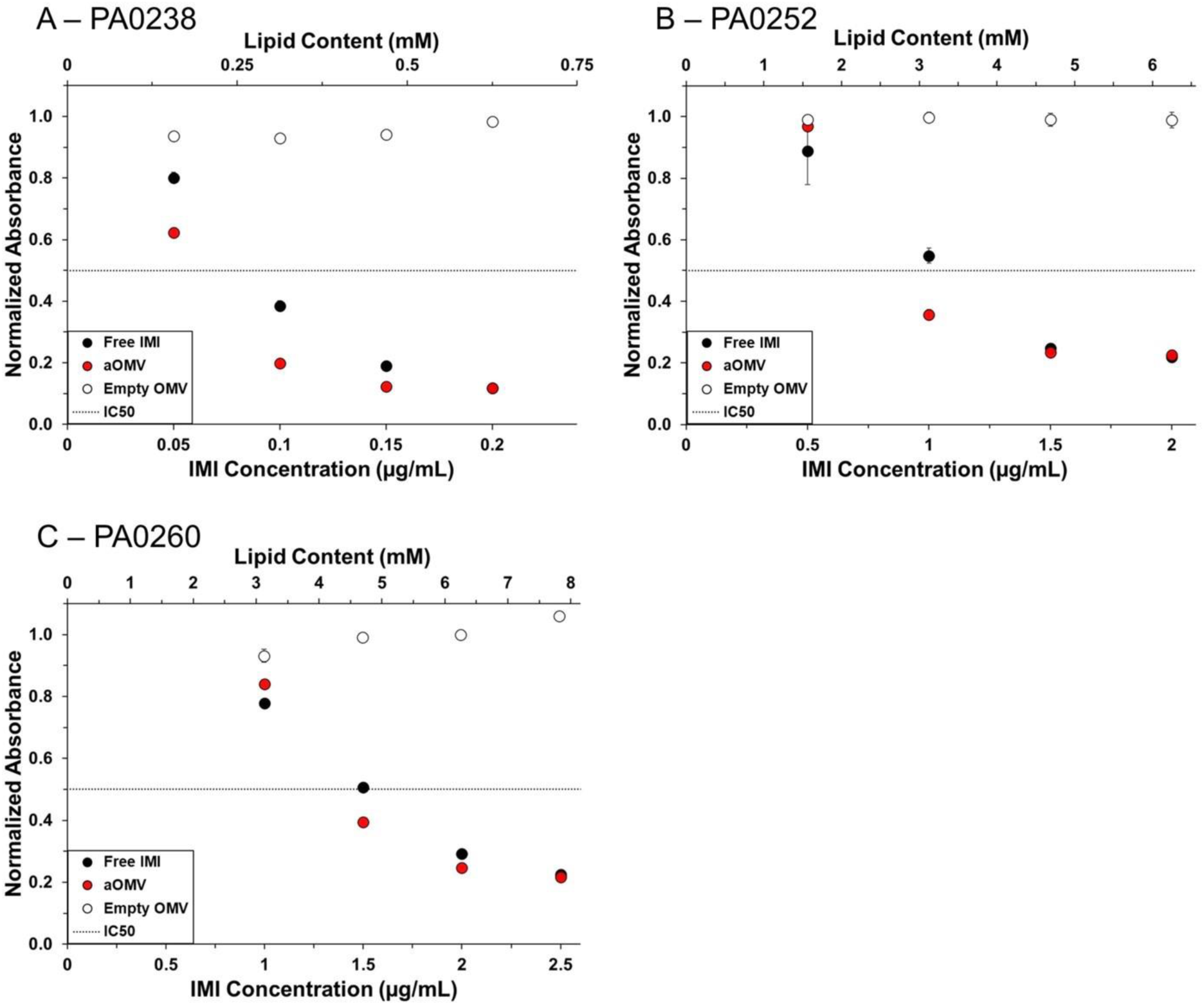
Delivery of IMI, both free and encapsulated within OMVs, to MDR *P. aeruginosa* clinical isolates (A) PA0238, (B) PA0252, and (C) PA0260. Bacteria were treated with free IMI (black), IMI-OMVs (red), or empty OMVs (white). The change in absorbance at 600 nm after 12 hr of growth of treated bacteria was normalized to that of untreated bacteria. Each data point represents the mean (n=3) ± standard deviation. The dashed line represents the IC50.

Isolate PA0238 was highly susceptible to IMI, as demonstrated by the IC50 of free IMI of 0.1 µg/mL (Fig. 3A, Fig. S2). IMI-OMVs, with an IMI concentration of 0.1 µg/mL, were even more effective in inhibiting bacterial growth than was the free drug. Empty OMVs at the same concentration had no effect on PA0238 growth, indicating that the enhanced effectiveness of IMI- OMVs was not due to any component of the OMVs themselves. Isolates PA0252 and PA0260 were less susceptible to IMI, with IC50 values of 1.5 and 2 µg/mL, respectively. IMI-OMVs improved the effectiveness of IMI in both isolates, lowering the IC50 values to 1.0 and 1.5 µg/mL respectively (Fig. 3B, 3C, S2). Again, these differences were not due to any components of the OMVs, as empty OMVs at the same lipid concentration did not inhibit bacterial growth. These results demonstrate that aOMVs can lower the inhibitory antibiotic concentration relative to free antibiotics in bacteria that are fully or moderately susceptible to the drug.

### 1. 3. Mechanism of aOMV delivery

To determine whether aOMVs enable transport of antibiotics in a porin-independent manner, as we hypothesized, we compared the growth of wild type *E. coli* K-12 BW25113, against several mutants from the Keio collection, including JW0912 (Δ*ompF*), JW2203 (Δ*ompC*), JW0940 (Δ*ompA*), and JW3368 (Δ*ompR*), each carrying a single-gene deletion^41^. OmpF is an essential pathway for the transport of small antibiotic drug molecules, including IMI across the membrane^42,43^. OmpA, on the other hand, has a low permeability for small ions and is mainly associated with membrane structural integrity^44,45^. OmpC has a role in both, maintaining structural integrity while allowing the passage of small molecules^42^. OmpR is a transcription factor that regulates the expression of porin genes in response to changes in the environmental osmotic pressure ^46,47^.

As demonstrated by the experimental IC50 of IMI against the parent strain and the four mutant strains (Table 2), the OmpF-deficient strain is less susceptible to the antibiotics compared to the wild type. The other mutant strains were either equally or more susceptible to the antibiotic. This IC50 data indicates that OmpF is essential for the delivery of IMI, while the OmpC, OmpA, and OmpR proteins do not play an important role in facilitating the transport of this antibiotic across the outer membrane.

**Table 2:**
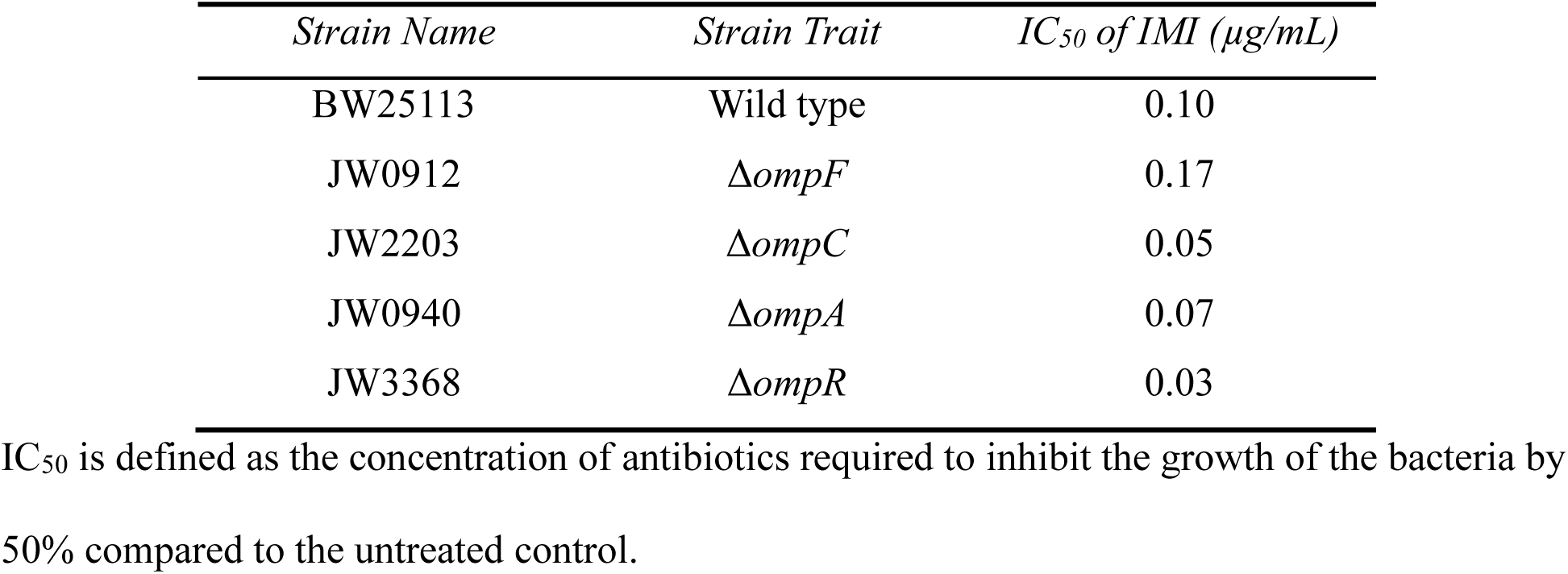
The experimental IC50 of IMI against *E. coli* Keio collection strains, reported in µg/mL.

| *Strain Name* | *Strain Trait* | *IC<sub>50</sub> of IMI ($\mu\text{g/mL}$)* |
| --- | --- | --- |
| BW25113 | Wild type | 0.10 |
| JW0912 | $\Delta ompF$ | 0.17 |
| JW2203 | $\Delta ompC$ | 0.05 |
| JW0940 | $\Delta ompA$ | 0.07 |
| JW3368 | $\Delta ompR$ | 0.03 |

IC<sub>50</sub> is defined as the concentration of antibiotics required to inhibit the growth of the bacteria by 50% compared to the untreated control.

To determine whether OMVs enable antibiotic delivery in the absence of outer membrane proteins, we grew the wild type and mutant strains in the presence of (1) empty OMVs, (2) free IMI, or (3) IMI-OMVs. In the absence of any treatment, all five strains grew in a similar manner (Fig. S3). Both IMI-OMVs and free IMI were equally effective in inhibiting the growth of the wild type strain (BW25113) at all concentrations (Fig. 4A, Fig. S4). In contrast, IMI-OMVs were significantly more effective than free IMI in treating JW0912 (Δ*ompF*) (Fig. 4B, Fig. S4). Empty OMVs did not contribute to the growth inhibition. For the other strains, there was no significant difference between the antibacterial potency of IMI-OMVs and free IMI, indicating that the absence of OmpC, OmpA or OmpR did not interfere with the internalization of IMI (Fig. 4C-E, Fig. S4). This result demonstrates that in the absence of OmpF, when free IMI is less effective due to its inability to cross the membrane, IMI-OMVs are able enhance the effectiveness of IMI by promoting its uptake.

**Figure 4:**
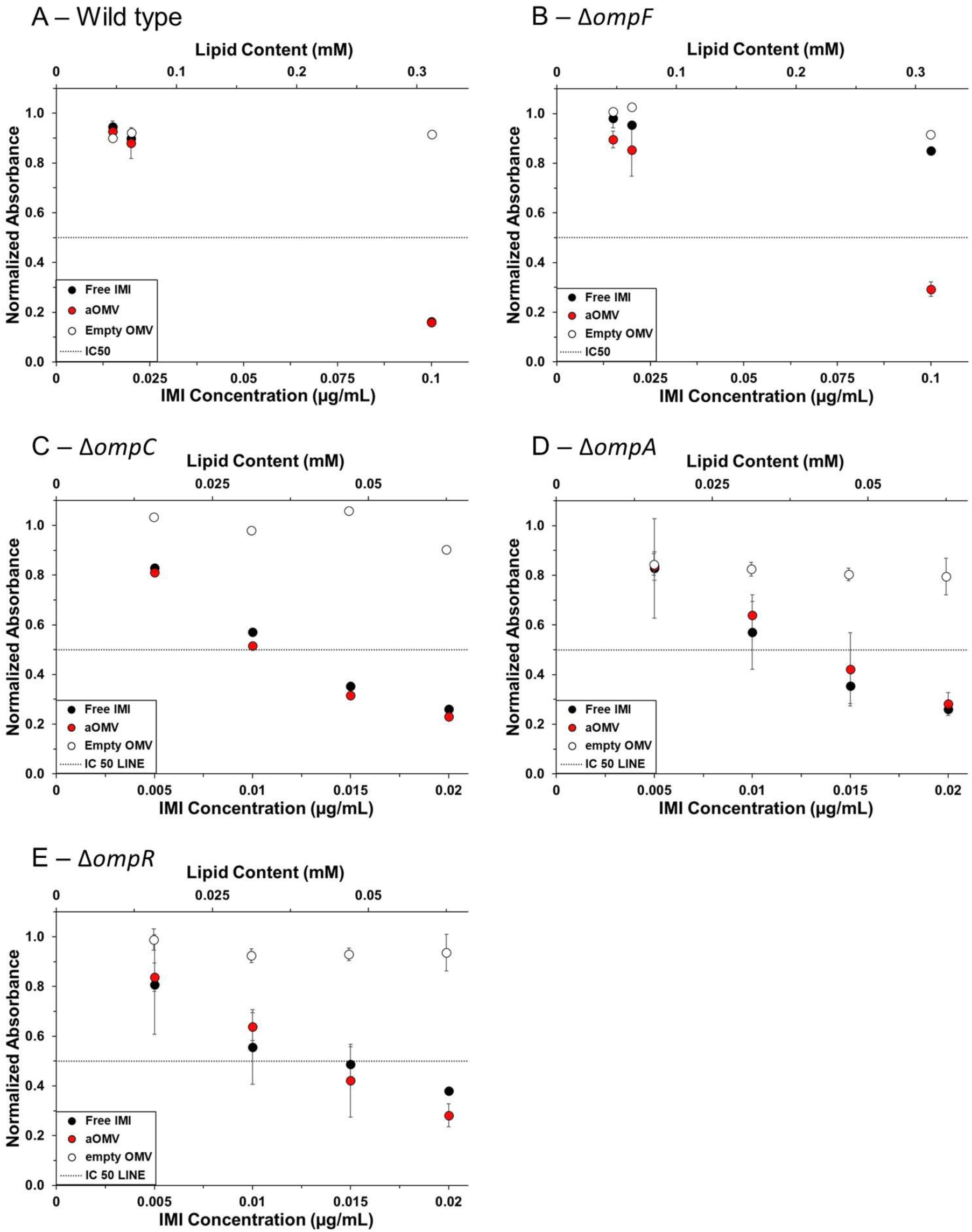
Treatment of *E. coli* Keio mutants with IMI. (A) BW25113 (wild type), (B) JW0912 (Δ*ompF*), (C) JW2203 (Δ*ompC*), (D) JW0940 (Δ*ompA*), and (E) JW3368 (Δ*ompR*). Each strain was treated with either free IMI (black), IMI-OMVs (red), or empty OMVs (white). The change in absorbance at 600 nm after 12 hr of growth of treated bacteria was normalized to that of untreated bacteria. The dashed line represents the IC50. Each data point represents the mean (n=3) ± standard deviation.

## 4. Delivery to Gram-positive bacteria

To determine whether aOMVs are effective against Gram-positive bacteria, we explored the effectiveness of aOMVs in treating borderline oxacillin-susceptible *S. aureus* (BORSA) clinical isolates from the CDC/FDA isolate bank. We observed that aOMVs were able to inhibit the growth of isolates SA0462 (Fig. 5A, Fig. S5) and SA0484 (Fig. 5B, Fig. S5). Empty OMVs did not contribute to the bactericidal activity.

**Figure 5:**
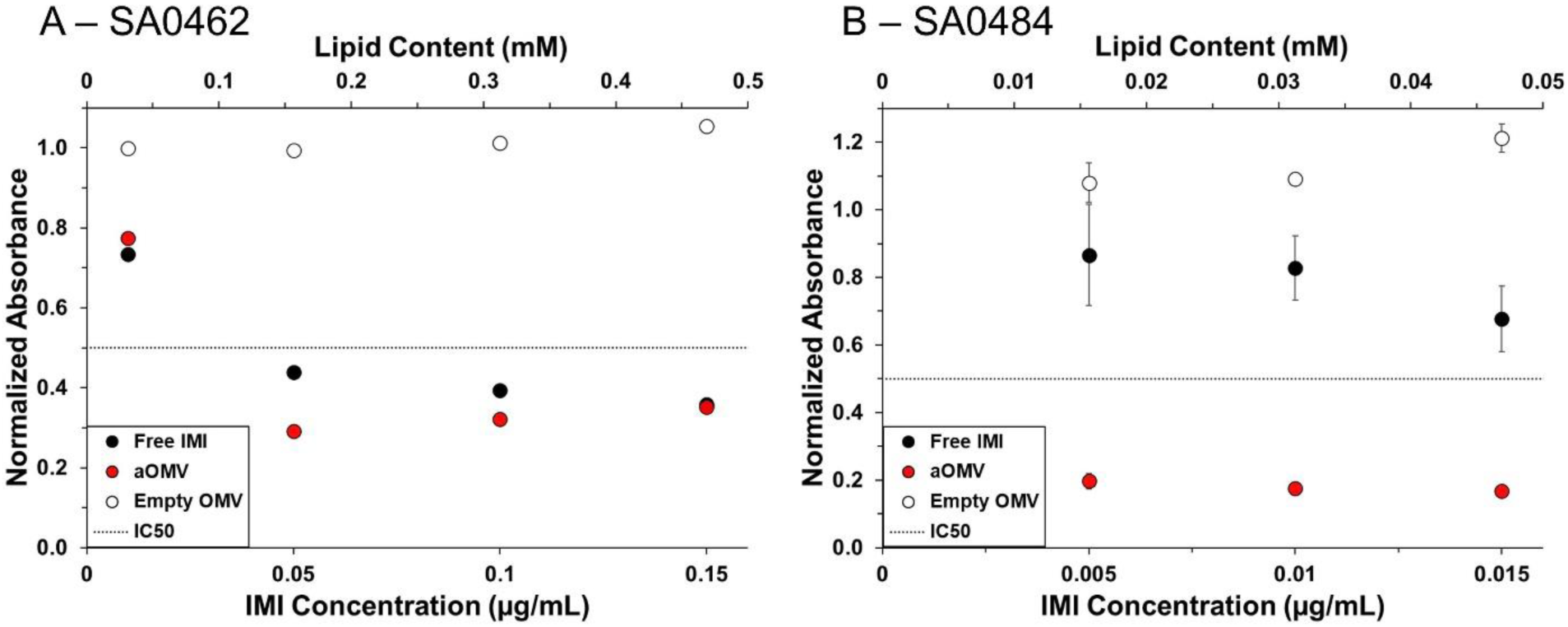
Delivery of IMI, both free and loaded, to BORSA clinical isolates, including (A) SA0462 and (B) SA0484. Each isolate was treated with either free IMI (black), IMI-OMVs (red), or empty OMVs (white). The dashed line represents the IC50. The change in absorbance at 600 nm after 12 hr of growth of treated bacteria was normalized to that of untreated bacteria. Each data point represents the mean (n=3) ± standard deviation.

## Discussion

In this study, we demonstrated that aOMVs can effectively inhibit the growth of both Gram-negative and Gram-positive bacteria, including antibiotic-resistant clinical isolates. Importantly, in several cases, encapsulation of the drugs within OMVs promoted the effectiveness of the antibiotics, as a lower concentration of antibiotics was needed to reach the same effect. While some OMVs have been reported to have bacteriolytic effects^48,49^, we showed that the *E. coli* JC8031 OMVs used in this study did not possess bacteriolytic activity. Thus, the increased effectiveness of the antibiotics is likely due to enhanced delivery across the Gram-negative membrane facilitated by the OMVs.

The outer membrane barrier limits the uptake of many antibiotics by Gram-negative bacteria^7,50^. The pathway of drug uptake by bacteria across the outer membrane is determined by the molecule’s hydrophobicity and charge^51,52^. Cationic aminoglycosides and polymyxins disrupt the outer membrane through electrostatic binding and gain access to the periplasmic space in the “self-promoted uptake” pathway^14,53–55^. Hydrophobic molecules can bind to the outer membrane and diffuse through the lipid bilayer^10,17,56^, while small hydrophilic molecules can only gain entry by passing through non-specific outer membrane proteins^10,57,58^. When resistant bacteria modify the outer membrane permeability and downregulate the expression of outer membrane porin channels, such as OmpF, the uptake of small hydrophilic molecules is limited^58–60^.

Multiple studies have demonstrated that OMVs are able to deliver their cargo across this barrier. This phenomenon has frequently been demonstrated by showing that OMVs collected from antibiotic-resistant organisms contain antibiotic resistance genes which can then be transferred to susceptible organisms. For example, clinical *Klebsiella pneumoniae* isolates were found to package the gene encoding for a specific carbapenemase (OXA-232) into OMVs, which were able to deliver those genes to non-resistant isolates^61^. A similar mechanism was observed to describe the horizontal transfer of the gene encoding for the OXA-24 β-lactamase by *Acinetobacter baumannii* OMVs^35^. Additionally, the hydrophobic nature of the OMV membrane enables the delivery of long-chain hydrophobic quorum sensing molecules, such as N-hexadecanoyl-L-homoserine lactone (C16-HSL) produced by *Paracoccus denitrificans*, and 2-heptyl-3-hydroxy- 4-quinolone (pseudomonas quinolone signal, PQS), produced by *P. aeruginosa* to other bacteria^62,63^. Although the detailed mechanisms of these OMV-mediated transport processes have not yet been identified, evidence suggests that OMVs are able to fuse with the outer membrane of Gram-negative bacteria. Specifically, Kadurugamuwa and Beveridge showed that after incubating *Salmonella typhimurium* or *E. coli* with OMVs produced by *Shigella flexneri* or *P. aeruginosa*, a significant amount of LPS from the donor bacteria could be detected on the surface of the acceptor bacteria, demonstrating the membrane mixing that occurs upon fusion^32,64^.

We therefore hypothesized that by encapsulating small hydrophilic antibiotics into OMVs, we could enable improved uptake of the antibiotics as a means of enhancing their activity. To investigate this hypothesis, we delivered IMI-OMVs to an OmpF-deficient mutant strain of *E. coli* and compared the resulting inhibition of growth against that of the wild type. We found that IMI- OMVs were more effective against the OmpF deficient strain than unencapsulated antibiotics at the same concentration. This finding supports our proposed mechanism by which aOMVs deliver luminal content via fusion with the bacterial outer membrane, thus delivering cargo in a porin-independent manner.

Our results build on the recent discovery by the Fuhrmann group demonstrating that OMVs from a myxobacterium (SBSr073) were effective in delivering ciprofloxacin to the enteropathogen, *Shigella flexneri*^65^. Thus, it appears that OMV-mediated delivery of luminal content may be a broad phenomenon among various Gram-negative bacterial species.

This observation that OMVs enable porin-independent delivery of antibiotics opens several exciting possibilities. There are many more antibiotics that are effective against Gram-positive bacteria than there are that are effective against Gram-negative bacteria. In many cases, these molecules have such a narrow spectrum of activity, simply because their physical properties do not allow them to cross the Gram-negative outer membrane^66^. For example, vancomycin, a tricyclic glycopeptide that irreversibly binds to D-alanyl-D-alanine moieties in peptidoglycan to disrupt the bacterial cell wall, resulting in cell lysis^67^ is too large to cross the outer membrane or to pass through porins. As a result, it is ineffective against Gram-negative bacteria because it is unable to access the peptidoglycan of these bacteria. We propose that OMVs could facilitate delivery of these types of molecules across the outer membrane, potentially increasing the effectiveness and broadening the spectrum of existing antibiotics, by enabling porin-independent delivery.

The mechanism by which the aOMVs interact with Gram-positive bacteria is less clear. However, a similar phenomenon has been reported previously. Kadurugamuwa and Beveridge showed that gentamicin-containing OMVs isolated from *P. aeruginosa* were active against *S. aureus* (strain D2C). The authors proposed that because of the high curvature of the OMVs, there is little salt bridging between adjacent LPS molecules (unlike on the bacterial surface); once the OMVs approach the peptidoglycan layer of the Gram-positive bacterium, which is enriched in divalent cations, the increased salt bridging leads to disruption of the OMV and release of encapsulated content, thus releasing a concentrated and localized dose of antibiotic to the cells^64^. Since then, several others have observed that fusogenic liposomes, which are designed to facilitate membrane fusion, increase the antibacterial activity against Gram-positive bacteria, including Staphylococci^22,68^. We hypothesize that a similar mechanism is responsible for the significant effectiveness of the OMVs against the *S. aureus* isolates observed here.

## Conclusions

In conclusion, this work demonstrates that aOMVs improve the activity of antibiotics by enhancing their transport across the Gram-negative bacterial membrane. This porin-independent delivery process enables improved activity of antibiotics, even when porin expression is decreased, and may be useful to allow delivery of large, narrow spectrum antibiotics to Gram-negative bacteria. We demonstrated that this delivery method is effective in Gram-negative and Gram-positive bacteria, in both lab strains and clinical antibiotic-resistant isolates. Current work is focused on elucidating the mechanism of OMV-outer membrane fusion to enable the design of biologically inspired synthetic systems with similar activity.

## Materials and Methods

1. Bacterial growth conditions

The list of bacteria used in these experiments is included in Table 3. All *P. aeruginosa* strains and *E. coli* strains, except for the Keio strains, were grown in Luria-Bertani (LB) broth Lennox from Invitrogen (Waltham, MA). The *E. coli* Keio strains, including the wild type and mutants, were cultivated in LB low salt broth, supplemented with 25 µg/mL of kanamycin. All clinical isolates of *S. aureus* were cultivated in tryptic soy broth from Becton Dickinson (Franklin Lakes, NJ).

1. 2. Chemicals

**Table 3:** List of bacteria used in this study.

| *Species* | *Strain Name* | *Notable Characteristics* | *Source* |
| --- | --- | --- | --- |
| *E. coli* | W3110 | K-12 wild type | 69 |
| | JC8031 | K-12 mutant, $\Delta tolRA$, hypervesiculating | 70 |
|  | BW25113 | Keio collection wild type | 41 |
| | JW0912 | Keio mutant, $\Delta ompF$ | 41 |
| | JW2203 | Keio mutant, $\Delta ompC$ | 41 |
| | JW0940 | Keio mutant, $\Delta ompA$ | 41 |
| | JW3368 | Keio mutant, $\Delta ompR$ | 41 |
| *P. aeruginosa* | PAO1 | Wild type | 71 |
|  | PA0238 | Clinical isolate from CDC | 72 |
|  | PA0252 | Clinical isolate from CDC | 72 |
|  | PA0260 | Clinical isolate from CDC | 72 |
| *S. aureus* | SA0462 | Clinical isolate from CDC | 72 |
|  | SA0484 | Clinical isolate from CDC | 72 |

IMI was purchased from Fisher Scientific (Waltham, MA). All chemicals were used without any additional purification.

1. 3. Isolation and purification of OMVs

A starter culture of *E. coli* JC8031 was made by combining 10 mL LB Lennox with 100 µL of a glycerol stock of the bacterium; this culture was grown at 37 °C with shaking at 175 RPM for 8 to 12 h. The starter culture was further diluted with LB at a 1:100 ratio and grown until the culture reached the late exponential phase. The supernatant was separated from the bacteria by centrifugation at 10,000 x g for 10 min at 4 °C, followed by a second centrifugation at 10,000 x g for 5 min at 4 °C in a new container. The supernatant was then filtered through a 0.45 µm polyethersulfone (PES) membrane to remove any remaining cells. Prior to ultracentrifugation, the supernatant was concentrated to approximately 150 mL using centrifugal concentrator tubes with a 50 kDa molecular weight cut-off (MWCO), by centrifuging at 5,000 x g for 10 min at 4 °C. The concentrated supernatant was then ultracentrifuged twice at 175,000 x g for 1 h at 4 °C. The pellet was resuspended in 1 mL phosphate buffered saline (PBS) and filtered through a 0.45 µm PES syringe filter before being stored at -20 °C for further use.

1. 4. Synthesis of aOMVs

Antibiotics were loaded into OMVs using sonication following a published protocol^40,73^ The OMVs were mixed with an equal volume of a 0.1% w/v antibiotic solution. The mixture was incubated at room temperature for 30 min, then sonicated in a bath sonicator (VWR) for 30 s at 35 kHz, then placed on ice for 60 s, followed by another sonication in the bath sonicator for 30 s at the same settings. The sonicated mixture was incubated at room temperature for 1 hr to allow OMV recovery. After loading was completed, the OMV-antibiotic mixture was transferred to an Amicon Ultra 30-kDa MWCO centrifugal filter and centrifuged at 14,000 x g for 15 min at 4 °C.

The filtrate, which contained unencapsulated antibiotics, was collected and diluted with PBS. The absorbance of each dilution was measured using a Tecan Infinite® 200 PRO plate reader. The EE was calculated using Equation 1, where 𝑎_𝑡𝑜𝑡𝑎𝑙_is the total mass of antibiotic added into the mixture; and 𝑎_𝑓𝑟𝑒𝑒_is the mass of unencapsulated antibiotic, which was determined by comparing the absorbance of the filtrate with a calibration curve (Fig. S6).

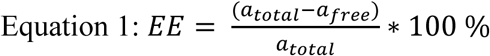

1. 5. Measurement of vesicle size

The diameter of the OMVs was measured using DLS. The sample was diluted with PBS at a 1:20 volume ratio and filtered through a PES membrane with 0.45 µm pore size to eliminate any potential contamination. We used an ALV/CGS-3 compact goniometer system, measured at a wavelength of 632.8 nm and scattering angle of 90°, with a run duration of 180 s. The data were collected in triplicate. The ALV built-in software was used for data processing. The number-weighted size distribution was plotted with the ALV built-in software program using a regularized fit and a membrane thickness of 5 nm (r* = 5 nm).

1. 6. Determination of lipid content

The lipid content of the OMVs was determined using the lipophilic FM^TM^ 4-64 dye (Invitrogen). Liposomes composed of 1-palmitoyl-2-oleoylglycero-3-phosphocholine (POPC) at known lipid concentrations were used as a comparison. After incubating 200 ng of the dye per 50µL of diluted sample in the dark for 15 s, the fluorescence spectra were recorded with an excitation wavelength of 515 nm and emission wavelength of 640 nm using a PTI QuantaMaster fluorometer.

1. 7. Scanning electron microscopy analysis

SEM was used to visualize the integrity and size of the OMVs. The images were collected using a Hitachi Scientific Instrument S4300SE Schottky-Emission SEM. The samples were first fixed using 2% glutaraldehyde for 2 hr in microcentrifuge tubes, then rapidly frozen in liquid nitrogen before being lyophilized. Following the deposition of lyophilized OMVs on aluminum stubs, the samples were coated with iridium using an EMS575X sputter coater, with a thickness of approximately 3 nm. Visualization was achieved using an accelerating voltage of 5 kV.

1. 8. aOMV effectiveness assay

The effectiveness of the aOMVs was analyzed by assessing their ability to inhibit bacterial growth over a range of antibiotic concentrations. The bacteria were treated with (1) PBS, (2) free antibiotics in a range of concentrations, (3) aOMVs with the same antibiotic concentrations as treatment 2, or (4) empty OMVs with the same lipid concentration as treatment 3. Bacterial growth at 37°C was monitored by collecting hourly readings of the OD600 using a Tecan Infinite® 200 PRO plate reader. All samples were prepared in triplicate in a sterile 96 well-plate. The data were normalized using the growth of untreated bacteria to compare the change in bacterial growth for each treatment.

1. 9. aOMV leakage

The stability of the aOMVs was assessed by determining the extent of leakage of encapsulated content under various conditions. aOMVs with known concentrations of encapsulated antibiotic were stored in a 37 °C incubator for 24 h. PBS was added to rinse any potential antibiotic from the aOMV surface and collect drug molecules leaked from the interior of aOMVs. The mixture was transferred to a concentrator with 30 kDa MWCO and centrifuged at 14,000 x g for 15 m to separate the leaked antibiotic and membrane fragments from intact aOMVs. The UV-Vis absorbance of the filtrate was compared against a calibration curve to calculate the concentration of released content.

### Supporting Information

Additional experimental results, including calibration curves and bacterial growth curves.

## Acknowledgment

This work was supported by Lehigh University (Faculty Innovation Grant (ACB) and Chen Fellowship (MW)).

We thank Dr. Matthew DeLisa for kindly contributing the *E. coli* JC8031 bacterial strain used in this study.

## References

(1) Centers for Disease Control, U. Antibiotic Resistance Threats in the United States, 2019. 2019. 10.15620/CDC:82532.

(2) Centers for Disease Control, U. COVID-19 : U.S. Impact on Antimicrobial Resistance, Special Report 2022. 2022. 10.15620/CDC:117915.

(3) Lewis, K. The Science of Antibiotic Discovery. Cell 2020, 181 (1), 29–45. 10.1016/J.CELL.2020.02.056.

(4) Boucher, H. W.; Talbot, G. H.; Bradley, J. S.; Edwards, J. E.; Gilbert, D.; Rice, L. B.; Scheld, M.; Spellberg, B.; Bartlett, J. Bad Bugs, No Drugs: No ESKAPE! An Update from the Infectious Diseases Society of America. Clinical Infectious Diseases 2009, 48 (1), 1–12. 10.1086/595011/2/48-1-1-TBL002.GIF.

(5) Beveridge, T. J. Structures of Gram-Negative Cell Walls and Their Derived Membrane Vesicles. J Bacteriol 1999, 181 (16), 4725–4733. 10.1128/JB.181.16.4725-4733.1999/ASSET/43C81F41-AF1C-4D3E-975B-854DA924259F/ASSETS/GRAPHIC/JB1590507009.JPEG.

(6) Costerton, J. W.; Cheng, K. J. The Role of the Bacterial Cell Envelope in Antibiotic Resistance. Journal of Antimicrobial Chemotherapy 1975, 1 (4), 363–377. 10.1093/JAC/1.4.363.

(7) Delcour, A. H. Outer Membrane Permeability and Antibiotic Resistance. Biochim Biophys Acta 2009, 1794 (5), 808. 10.1016/J.BBAPAP.2008.11.005.

(8) Nikaido, H. Molecular Basis of Bacterial Outer Membrane Permeability Revisited. Microbiology and Molecular Biology Reviews 2003, 67 (4), 593–656. 10.1128/mmbr.67.4.593-656.2003.

(9) Webber, M. A.; Piddock, L. J. V. The Importance of Efflux Pumps in Bacterial Antibiotic Resistance. Journal of Antimicrobial Chemotherapy 2003, 51 (1), 9–11. 10.1093/JAC/DKG050.

(10) Pagès, J.-M.; James, C. E.; Winterhalter, M. The Porin and the Permeating Antibiotic: A Selective Diffusion Barrier in Gram-Negative Bacteria. 2008. 10.1038/nrmicro1994.

(11) O. Gutkind, G.; Di Conza, J.; Power, P.; Radice, M. β-Lactamase-Mediated Resistance: A Biochemical, Epidemiological and Genetic Overview. Curr Pharm Des 2013, 19 (2), 164–208. 10.2174/1381612811306020164.

(12) Aldred, K. J.; Kerns, R. J.; Osheroff, N. Mechanism of Quinolone Action and Resistance. Biochemistry 2014, 53 (10), 1565. 10.1021/BI5000564.

(13) Wanger, A.; Chavez, V.; Huang, R. S. P.; Wahed, A.; Actor, J. K.; Dasgupta, A. Antibiotics, Antimicrobial Resistance, Antibiotic Susceptibility Testing, and Therapeutic Drug Monitoring for Selected Drugs. Microbiology and Molecular Diagnosis in Pathology 2017, 119–153. 10.1016/B978-0-12-805351-5.00007-7.

(14) Mohapatra, S. S.; Dwibedy, S. K.; Padhy, I. Polymyxins, the Last-Resort Antibiotics: Mode of Action, Resistance Emergence, and Potential Solutions. J Biosci 2021, 46 (3). 10.1007/S12038-021-00209-8.

(15) Prachayasittikul, V.; Isarankura-Na-Ayudhya, C.; Tantimongcolwat, T.; Nantasenamat, C.; Galla, H. J. EDTA-Induced Membrane Fluidization and Destabilization: Biophysical Studies on Artificial Lipid Membranes. Acta Biochim Biophys Sin (Shanghai) 2007, 39 (11), 901–913. 10.1111/J.1745-7270.2007.00350.X.

(16) Vaara, M. Agents That Increase the Permeability of the Outer Membrane. Microbiol Rev 1992, 56 (3), 395. 10.1128/MR.56.3.395-411.1992.

(17) Flaten, G. E.; Dhanikula, A. B.; Luthman, K.; Brandl, M. Drug Permeability across a Phospholipid Vesicle Based Barrier: A Novel Approach for Studying Passive Diffusion. European Journal of Pharmaceutical Sciences 2006, 27 (1), 80–90. 10.1016/J.EJPS.2005.08.007.

(18) Li, J.; Koh, J. J.; Liu, S.; Lakshminarayanan, R.; Verma, C. S.; Beuerman, R. W. Membrane Active Antimicrobial Peptides: Translating Mechanistic Insights to Design. Front Neurosci 2017, 11 (FEB), 73. 10.3389/FNINS.2017.00073.

(19) Heindorff, K.; Aurich, O.; Michaelis, A.; Rieger, R. Genetic Toxicology of Ethylenediaminetetraacetic Acid (EDTA). Mutat Res 1983, 115 (2), 149–173. 10.1016/0165-1110(83)90001-5.

(20) Csiszár, A.; Hersch, N.; Dieluweit, S.; Biehl, R.; Merkel, R.; Hoffmann, B. Novel Fusogenic Liposomes for Fluorescent Cell Labeling and Membrane Modification. Bioconjug Chem 2010, 21 (3), 537–543. 10.1021/BC900470Y/ASSET/IMAGES/LARGE/BC-2009-00470Y_0007.JPEG.

(21) Kube, S.; Hersch, N.; Naumovska, E.; Gensch, T.; Hendriks, J.; Franzen, A.; Landvogt, L.;Siebrasse, J. P.; Kubitscheck, U.; Hoffmann, B.; Merkel, R.; Csiszár, A. Fusogenic Liposomes as Nanocarriers for the Delivery of Intracellular Proteins. Langmuir 2017, 33 (4), 1051–1059. 10.1021/ACS.LANGMUIR.6B04304/ASSET/IMAGES/LARGE/LA-2016-04304K_0003.JPEG.

(22) Nicolosi, D.; Cupri, S.; Genovese, C.; Tempera, G.; Mattina, R.; Pignatello, R. Nanotechnology Approaches for Antibacterial Drug Delivery: Preparation and Microbiological Evaluation of Fusogenic Liposomes Carrying Fusidic Acid. Int J Antimicrob Agents 2015, 45 (6), 622–626. 10.1016/J.IJANTIMICAG.2015.01.016.

(23) Sharma, A.; Gupta, V. K.; Pathania, R. Efflux Pump Inhibitors for Bacterial Pathogens: From Bench to Bedside. Indian J Med Res 2019, 149 (2), 129. 10.4103/IJMR.IJMR_2079_17.

(24) Liu, P.; Chen, G.; Zhang, J. A Review of Liposomes as a Drug Delivery System: Current Status of Approved Products, Regulatory Environments, and Future Perspectives. Molecules 2022, 27 (4). 10.3390/MOLECULES27041372.

(25) Schwechheimer, C.; Kuehn, M. J. Outer-Membrane Vesicles from Gram-Negative Bacteria: Biogenesis and Functions. Nature Reviews Microbiology 2015 13:10 2015, 13 (10), 605–619. 10.1038/nrmicro3525.

(26) Kulp, A.; Kuehn, M. J. Biological Functions and Biogenesis of Secreted Bacterial Outer Membrane Vesicles. Annual Review of Microbiology. October 13, 2010, pp 163–184. 10.1146/annurev.micro.091208.073413.

(27) Renelli, M.; Matias, V.; Lo, R. Y.; Beveridge, T. J. DNA-Containing Membrane Vesicles of Pseudomonas Aeruginosa PAO1 and Their Genetic Transformation Potential. Microbiology (N Y) 2004, 150 (7), 2161–2169. 10.1099/MIC.0.26841-0/CITE/REFWORKS.

(28) Dauros-Singorenko, P.; Blenkiron, C.; Phillips, A.; Swift, S. The Functional RNA Cargo of Bacterial Membrane Vesicles. FEMS Microbiol Lett 2018, 365 (5), 23. 10.1093/FEMSLE/FNY023.

(29) Brameyer, S.; Plener, L.; Müller, A.; Klingl, A.; Wanner, G.; Jung, K. Outer Membrane Vesicles Facilitate Trafficking of the Hydrophobic Signaling Molecule CAI-1 between Vibrio Harveyi Cells. J Bacteriol 2018, 200 (15). 10.1128/JB.00740-17/SUPPL_FILE/ZJB999094745S1.PDF.

(30) Ciofu, O.; Beveridge, T. J.; Kadurugamuwa, J.; Walther-Rasmussen, J.; Høiby, N. Chromosomal Beta-Lactamase Is Packaged into Membrane Vesicles and Secreted from Pseudomonas Aeruginosa. J Antimicrob Chemother 2000, 45 (1), 9–13. 10.1093/JAC/45.1.9.

(31) Chatterjee, D.; Chaudhuri, K. Association of Cholera Toxin with Vibrio Cholerae Outer Membrane Vesicles Which Are Internalized by Human Intestinal Epithelial Cells. FEBS Lett 2011, 585 (9), 1357–1362. 10.1016/J.FEBSLET.2011.04.017.

(32) Kadurugamuwat, J. L.; Beveridge, T. J. Membrane Vesicles Derived from Pseudornonas. Aeruginosa and Shigella Flexneri Can Be Integrated into the Surfaces of Other Gram-Negative Bacteria; 1999; Vol. 145.

(33) Lee, A. R.; Park, S. Bin; Kim, S. W.; Jung, J. W.; Chun, J. H.; Kim, J.; Kim, Y. R.; Lazarte, J. M. S.;Jang, H. Bin; Thompson, K. D.; Jung, M.; Ha, M. W.; Jung, T. S. Membrane Vesicles from Antibiotic-resistant Staphylococcus Aureus Transfer Antibiotic-resistance to Antibiotic-susceptible Escherichia Coli. J Appl Microbiol 2022, 132 (4), 2746–2759. 10.1111/JAM.15449.

(34) Shen, Z.; Qin, J.; Xiang, G.; Chen, T.; Nurxat, N.; Gao, Q.; Wang, C.; Zhang, H.; Liu, Y.; Li, M. Outer Membrane Vesicles Mediating Horizontal Transfer of the Epidemic BlaOXA-232 Carbapenemase Gene among Enterobacterales. Emerg Microbes Infect 2024, 13 (1). 10.1080/22221751.2023.2290840.

(35) Rumbo, C.; Fernández-Moreira, E.; Merino, M.; Poza, M.; Mendez, J. A.; Soares, N. C.; Mosquera, A.; Chaves, F.; Bou, G. Horizontal Transfer of the OXA-24 Carbapenemase Gene via Outer Membrane Vesicles: A New Mechanism of Dissemination of Carbapenem Resistance Genes in Acinetobacter Baumannii. Antimicrob Agents Chemother 2011, 55 (7), 3084–3090. 10.1128/AAC.00929-10/ASSET/45B25A80-FFC1-4B91-8B7D-81B1AA375388/ASSETS/GRAPHIC/ZAC9991000070005.JPEG.

(36) Chatterjee, S.; Mondal, A.; Mitra, S.; Basu, S. Acinetobacter Baumannii Transfers the BlaNDM-1 Gene via Outer Membrane Vesicles. Journal of Antimicrobial Chemotherapy 2017, 72 (8), 2201–2207. 10.1093/JAC/DKX131.

(37) Dorward, D. W.; Garon, C. F.; Judd, R. C. Export and Intercellular Transfer of DNA via Membrane Blebs of Neisseria Gonorrhoeae. J Bacteriol 1989, 171 (5), 2499–2505. 10.1128/JB.171.5.2499-2505.1989.

(38) Alves, N. J.; Turner, K. B.; Medintz, I. L.; Walper, S. A. Protecting Enzymatic Function through Directed Packaging into Bacterial Outer Membrane Vesicles. Scientific Reports 2016 6:1 2016, 6 (1), 1–10. 10.1038/srep24866.

(39) Alves, N. J.; Turner, K. B.; Daniele, M. A.; Oh, E.; Medintz, I. L.; Walper, S. A. Bacterial Nanobioreactors-Directing Enzyme Packaging into Bacterial Outer Membrane Vesicles. ACS Appl Mater Interfaces 2015, 7 (44), 24963–24972. 10.1021/ACSAMI.5B08811/ASSET/IMAGES/LARGE/AM-2015-08811Q_0008.JPEG.

(40) Wu, M.; Holgado, L.; Harrower, R. M.; Brown, A. C. Evaluation of the Efficiency of Various Methods to Load Fluoroquinolones into Escherichia Coli Outer Membrane Vesicles as a Novel Antibiotic Delivery Platform. Biochem Eng J 2024, 210. 10.1016/J.BEJ.2024.109418.

(41) Baba, T.; Ara, T.; Hasegawa, M.; Takai, Y.; Okumura, Y.; Baba, M.; Datsenko, K. A.; Tomita, M.; Wanner, B. L.; Mori, H. Construction of Escherichia Coli K-12 in-Frame, Single-Gene Knockout Mutants: The Keio Collection. Mol Syst Biol 2006, 2, 2006.0008. 10.1038/MSB4100050.

(42) Masi, M.; Pagès, J.-M. Structure, Function and Regulation of Outer Membrane Proteins Involved in Drug Transport in Enterobactericeae: The OmpF/C – TolC Case. Open Microbiol J 2013, 7 (1), 22. 10.2174/1874285801307010022.

(43) Kishii, R.; Takei, M. Relationship between the Expression of OmpF and Quinolone Resistance in Escherichia Coli. J Infect Chemother 2009, 15 (6), 361–366. 10.1007/S10156-009-0716-6.

(44) Wang, X.; Bernstein, H. D. The Escherichia Coli Outer Membrane Protein OmpA Acquires Secondary Structure Prior to Its Integration into the Membrane. Journal of Biological Chemistry 2022, 298 (4), 101802. 10.1016/J.JBC.2022.101802.

(45) Sugawara, E.; Nikaido, H. Pore-Forming Activity of OmpA Protein of Escherichia Coli. Journal of Biological Chemistry 1992, 267 (4), 2507–2511. 10.1016/S0021-9258(18)45908-X.

(46) Chakraborty, S.; Kenney, L. J. A New Role of OmpR in Acid and Osmotic Stress in Salmonella and E. Coli. Front Microbiol 2018, 9 (NOV). 10.3389/fmicb.2018.02656.

(47) Gerken, H.; Vuong, P.; Soparkar, K.; Misra, R. Roles of the ENVZ/OMPR Two-Component System and Porins in Iron Acquisition in Escherichia Coli. mBio 2020, 11 (3), 1–18. 10.1128/MBIO.01192-20/SUPPL_FILE/MBIO.01192-20-SF006.TIF.

(48) Kadurugamuwa, J. L.; Beveridge, T. J. Membrane Vesicles Derived from Pseudomonas Aeruginosa and Shigella Flexneri Can Be Integrated into the Surfaces of Other Gram-Negative Bacteria. Microbiology (N Y) 1999, 145 (8), 2051–2060. 10.1099/13500872-145-8-2051/CITE/REFWORKS.

(49) Kadurugamuwa, J. L.; Beveridge, T. J. Bacteriolytic Effect of Membrane Vesicles from Pseudomonas Aeruginosa on Other Bacteria Including Pathogens: Conceptually New Antibiotics. J Bacteriol 1996, 178 (10), 2767–2774. 10.1128/JB.178.10.2767-2774.1996.

(50) Blair, J. M.; Piddock, L. J.; Walsh, C.; Wright, G. Structure, Function and Inhibition of RND Efflux Pumps in Gram-Negative Bacteria: An Update This Review Comes from a Themed Issue on Antimicrobials Edited By. Curr Opin Microbiol 2009, 12, 512–519. 10.1016/j.mib.2009.07.003.

(51) Yang, N. J.; Hinner, M. J. Getting Across the Cell Membrane: An Overview for Small Molecules, Peptides, and Proteins. Methods Mol Biol 2015, 1266, 29. 10.1007/978-1-4939-2272-7_3.

(52) Zhang, R.; Qin, X.; Kong, F.; Chen, P.; Pan, G. Improving Cellular Uptake of Therapeutic Entities through Interaction with Components of Cell Membrane. Drug Deliv 2019, 26 (1), 328. 10.1080/10717544.2019.1582730.

(53) Nakae, R.; Nakae, T. Diffusion of Aminoglycoside Antibiotics Across the Outer Membrane of Escherichia Coli; 1982; Vol. 22.

(54) Krause, K. M.; Serio, A. W.; Kane, T. R.; Connolly, L. E. Aminoglycosides: An Overview. Cold Spring Harb Perspect Med 2016, 6 (6). 10.1101/CSHPERSPECT.A027029.

(55) Conceição-Neto, O. C.; da Costa, B. S.; Pontes, L. da S.; Silveira, M. C.; Justo-da-Silva, L. H.; de Oliveira Santos, I. C.; Teixeira, C. B. T.; Tavares e Oliveira, T. R.; Hermes, F. S.; Galvão, T. C.; Antunes, L. C. M.; Rocha-de-Souza, C. M.; Carvalho-Assef, A. P. D. Polymyxin Resistance in Clinical Isolates of K. Pneumoniae in Brazil: Update on Molecular Mechanisms, Clonal Dissemination and Relationship With KPC-Producing Strains. Front Cell Infect Microbiol 2022, 12, 898125. 10.3389/FCIMB.2022.898125/BIBTEX.

(56) Wardhan, R.; Mudgal, P. Membrane Transport. Textbook of Membrane Biology 2017, 149. 10.1007/978-981-10-7101-0_6.

(57) Acosta-Gutiérrez, S.; Ferrara, L.; Pathania, M.; Masi, M.; Wang, J.; Bodrenko, I.; Zahn, M.; Winterhalter, M.; Stavenger, R. A.; Pagès, J.-M.; Naismith, J. H.; van den Berg, B.; P Page, M. G.; Ceccarelli, M. Getting Drugs into Gram-Negative Bacteria: Rational Rules for Permeation through General Porins.

(58) Choi, U.; Lee, C. R. Distinct Roles of Outer Membrane Porins in Antibiotic Resistance and Membrane Integrity in Escherichia Coli. Front Microbiol 2019, 10 (APR), 953. 10.3389/FMICB.2019.00953/BIBTEX.

(59) Novikova, O. D.; Solovyeva, T. F. Nonspecific Porins of the Outer Membrane of Gram-Negative Bacteria: Structure and Functions. Biochem (Mosc) Suppl Ser A Membr Cell Biol 2009, 3 (1), 3–15. 10.1134/S1990747809010024/METRICS.

(60) Zhou, G.; Wang, Q.; Wang, Y.; Wen, X.; Peng, H.; Peng, R.; Shi, Q.; Xie, X.; Li, L. Outer Membrane Porins Contribute to Antimicrobial Resistance in Gram-Negative Bacteria. Microorganisms 2023, 11 (7). 10.3390/MICROORGANISMS11071690.

(61) Shen, Z.; Qin, J.; Xiang, G.; Chen, T.; Nurxat, N.; Gao, Q.; Wang, C.; Zhang, H.; Liu, Y.; Li, M. Outer Membrane Vesicles Mediating Horizontal Transfer of the Epidemic BlaOXA-232 Carbapenemase Gene among Enterobacterales. Emerg Microbes Infect 2024, 13 (1). 10.1080/22221751.2023.2290840/SUPPL_FILE/TEMI_A_2290840_SM6078.D OCX.

(62) Toyofuku, M.; Morinaga, K.; Hashimoto, Y.; Uhl, J.; Shimamura, H.; Inaba, H.; Schmitt-Kopplin, P.; Eberl, L.; Nomura, N. Membrane Vesicle-Mediated Bacterial Communication. ISME J 2017, 11 (6), 1504–1509. 10.1038/ISMEJ.2017.13.

(63) Mashburn, L. M.; Whiteley, M. Membrane Vesicles Traffic Signals and Facilitate Group Activities in a Prokaryote. Nature 2005 437:7057 2005, 437 (7057), 422–425. 10.1038/nature03925.

(64) Kadurugamuwa, J. L.; Beveridge, T. J. Bacteriolytic Effect of Membrane Vesicles from Pseudomonas Aeruginosa on Other Bacteria Including Pathogens: Conceptually New Antibiotics. J Bacteriol 1996, 178 (10), 2767–2774. 10.1128/JB.178.10.2767-2774.1996.

(65) Heinrich, E.; Hartwig, O.; Walt, C.; Kardani, A.; Koch, M.; Jahromi, L. P.; Hoppstädter, J.; Kiemer, A. K.; Loretz, B.; Lehr, C. M.; Fuhrmann, G. Cell-Derived Vesicles for Antibiotic Delivery—Understanding the Challenges of a Biogenic Carrier System. Small 2023. 10.1002/smll.202207479.

(66) O’Shea, R.; Moser, H. E. Physicochemical Properties of Antibacterial Compounds: Implications for Drug Discovery. J Med Chem 2008, 51 (10), 2871–2878. 10.1021/JM700967E/SUPPL_FILE/JM700967E.XLS.

(67) Sheldrick, G. M.; Jones, P. G.; Kennard, O.; Williams, D. H.; Smith, G. A. Structure of Vancomycin and Its Complex with Acetyl-D-Alanyl-D-Alanine. Nature 1978, 271 (5642), 223–225. 10.1038/271223A0.

(68) Scriboni, A. B.; Couto, V. M.; De Morais Ribeiro, L. N.; Freires, I. A.; Groppo, F. C.; De Paula, E.; Franz-Montan, M.; Cogo-Müller, K. Fusogenic Liposomes Increase the Antimicrobial Activity of Vancomycin Against Staphylococcus Aureus Biofilm. Front Pharmacol 2019, 10. 10.3389/FPHAR.2019.01401.

(69) Bachmann, B. J. Pedigrees of Some Mutant Strains of Escherichia Coli K-12. Bacteriol Rev 1972, 36 (4), 525. 10.1128/BR.36.4.525-557.1972.

(70) Bernadac, A.; Gavioli, M.; Lazzaroni, J. C.; Raina, S.; Lloubès, R. Escherichia Coli Tol-Pal Mutants Form Outer Membrane Vesicles. J Bacteriol 1998, 180 (18), 4872–4878. 10.1128/JB.180.18.4872-4878.1998/FORMAT/EPUB.

(71) Holloway, B. W. Genetic Recombination in Pseudomonas Aeruginosa. J Gen Microbiol 1955, 13 (3), 572–581. 10.1099/00221287-13-3-572/CITE/REFWORKS.

72. (72) Centers for Disease Control and Prevention, U. *FDA & CDC Antimicrobial Resistance Isolate Bank*. https://wwwn.cdc.gov/ARIsolateBank/ (accessed 2024-04-03).

(73) Lamichhane, T. N.; Jeyaram, A.; Patel, D. B.; Parajuli, B.; Livingston, N. K.; Arumugasaamy, N.; Schardt, J. S.; Jay, S. M. Oncogene Knockdown via Active Loading of Small RNAs into Extracellular Vesicles by Sonication. Cell Mol Bioeng 2016, 9 (3), 315–324. 10.1007/S12195-016-0457-4.

