## Supporting Information for "*Escherichia coli* outer membrane vesicles encapsulating small molecule antibiotics improve drug function by facilitating transport"

[1]Department of Chemical and Biomolecular Engineering, Lehigh University, 124 E. Morton St., Bethlehem, PA, 18015, USA

[2]Department of Biological Sciences, Lehigh University, 111 Research Dr., Bethlehem, PA, 18015, USA

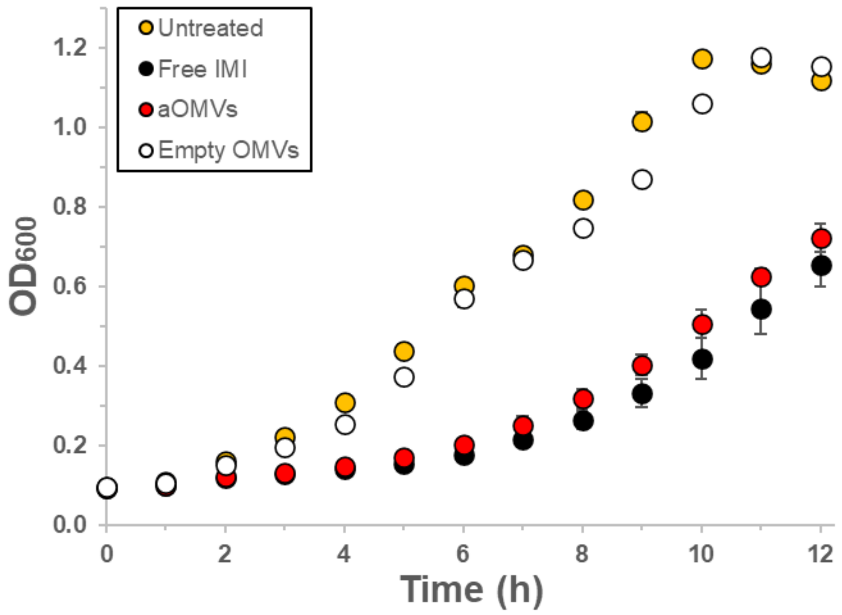


**Figure S1: Representative growth curves for *P. aeruginosa* PAO1.** The bacteria were untreated (yellow) or treated with free imipenem (black), aOMVs (red), or empty OMVs (white). The IMI concentration was 0.5 µg/mL. Each data point represents the mean (n=3) ± standard deviation.

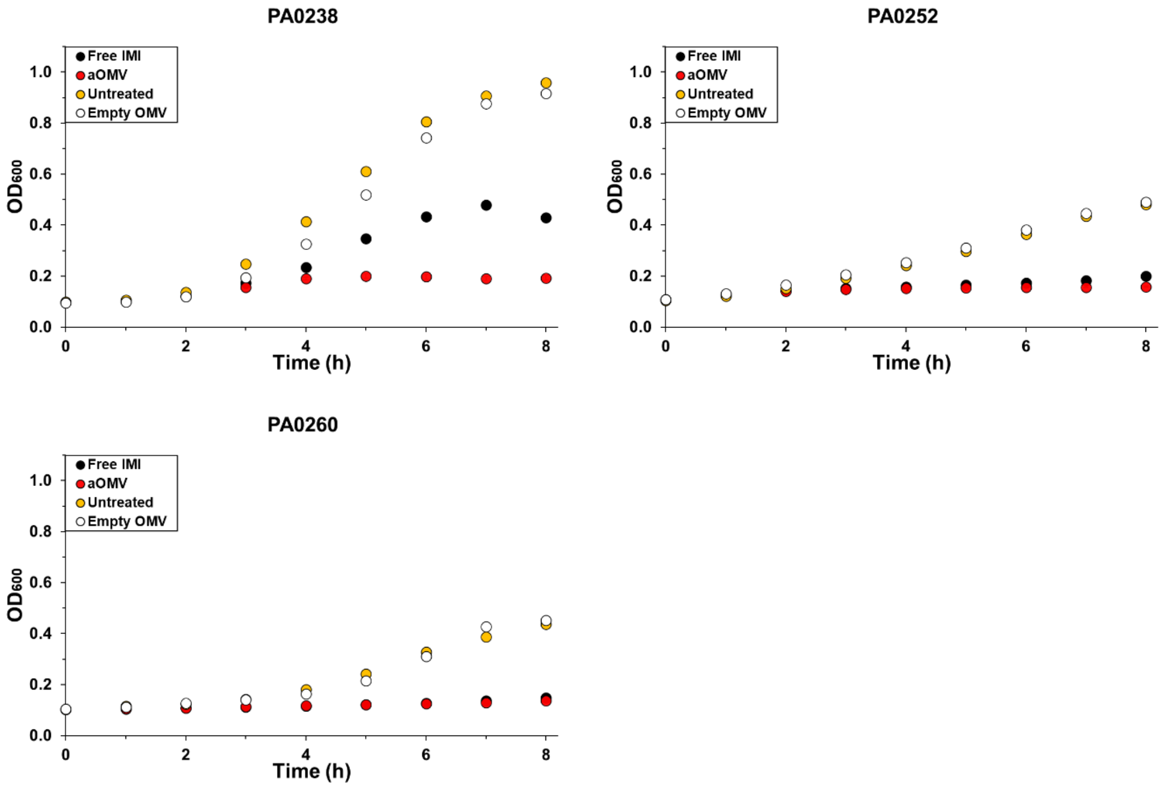


**Figure S2: Representative growth curves for (A) PA0238, (B) PA0252, and (C) PA0260.** The bacteria were untreated (yellow) or treated with free imipenem (black), aOMVs (red), or empty OMVs (white). PA0238 was treated with 0.1 µg/mL imipenem (free or in aOMVs), PA0252 was treated with 1 µg/mL imipenem (free or in aOMVs), and PA0260 was treated with 1.5 µg/mL imipenem (free or in aOMVs). Each data point represents the mean (n=3) ± standard deviation.

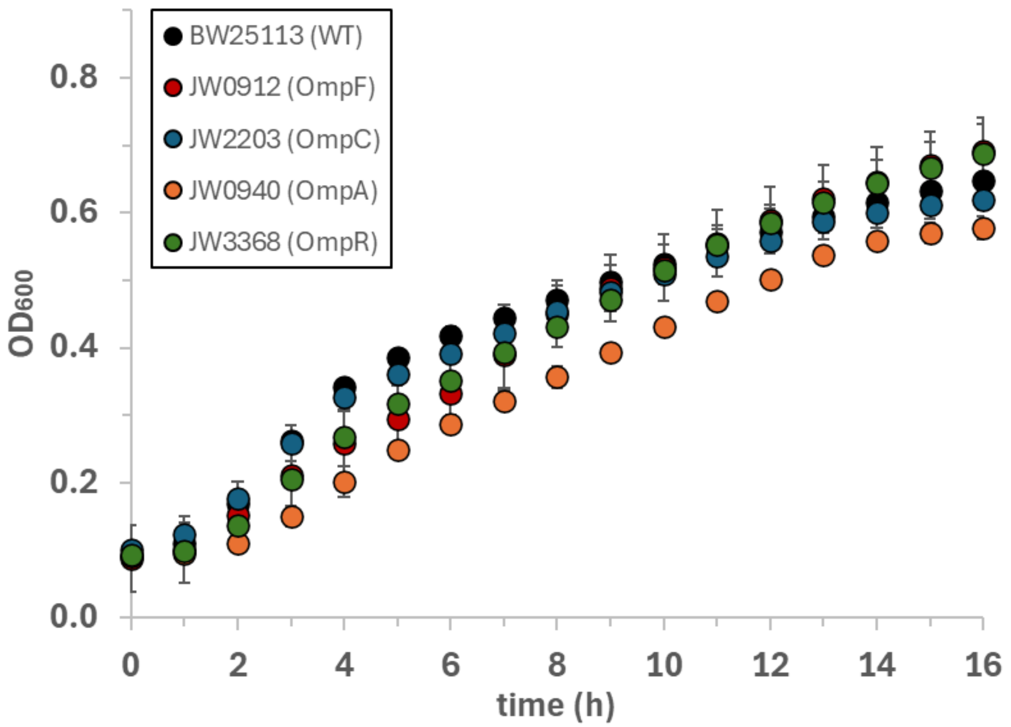


**Figure S3: Representative growth curves of Keio mutants.** The strains used in this work included BW25113 (wildtype, black), JS0912 (Δ*ompF*, red), JW2203 (Δ*ompC*, blue), JW0940 (Δ*ompA*, orange), and JW3368 (Δ*ompR*, green). All strains were grown in LB low salt broth, supplemented with 25 μg/mL of kanamycin. Each data point represents the mean (n=3) ± standard deviation.

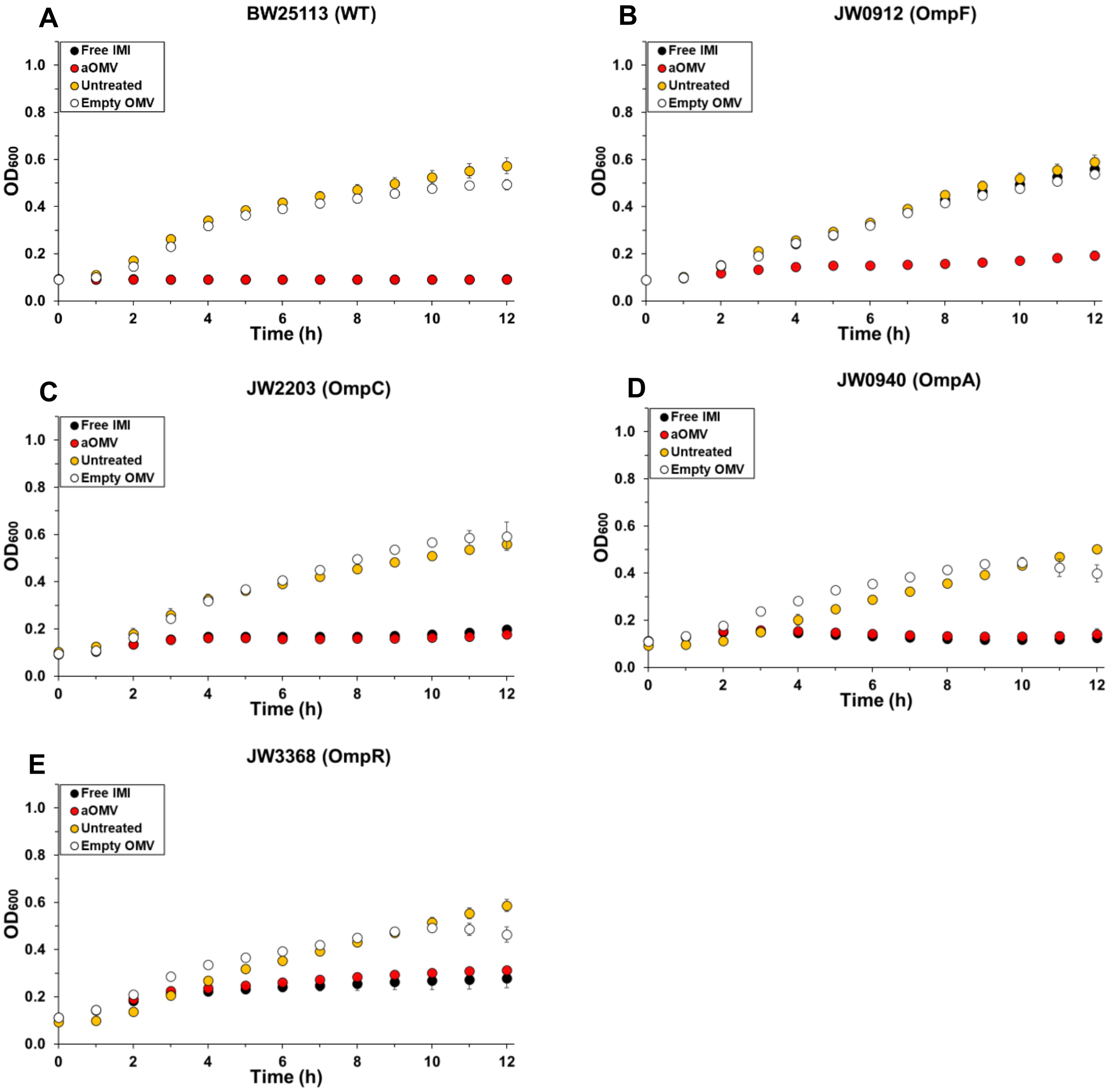


**Figure S4: Representative growth curves for Keio mutants treated with IMI.** (A) BW25113 (wildtype), (B) JW0912 (Δ*ompF*), (C) JW2203 (*ΔompC*), (D) JW0940 (*ΔompA*), and JW3368 (*ΔompR*). The bacteria were untreated (yellow) or treated with free IMI (0.01 µg/mL, black), aOMVs with an IMI concentration of 0.01 µg/mL (red), or empty OMVs with the same lipid concentration as the aOMVs (white). Each data point represents the mean (n=3) ± standard deviation.

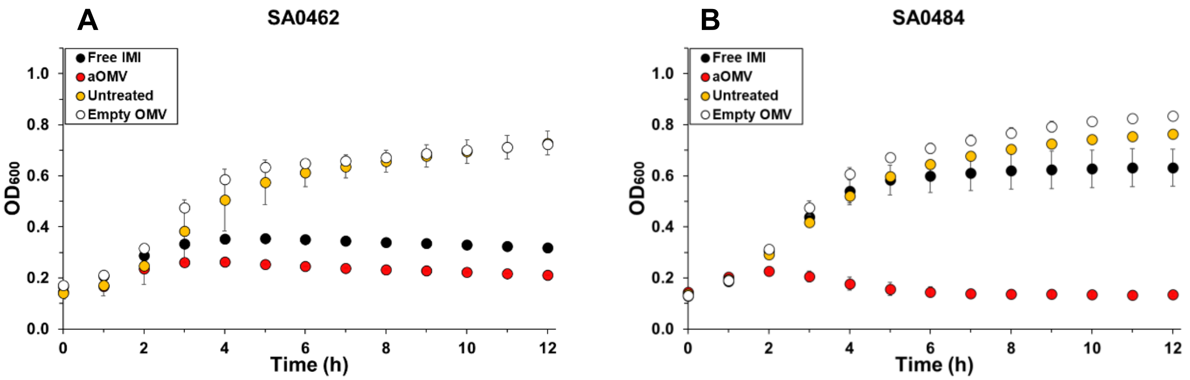


**Figure S5: Representative growth curves for BORSA clinical isolates (A) SA0462 and (B) SA0484**. (A) Isolate SA0462 was untreated (yellow) or treated with free IMI (0.05 µg/mL, black), aOMVs with an IMI concentration of 0.05 µg/mL (red), or empty OMVs with the same lipid concentration as the aOMVs (white). (B) Isolate SA0484 was untreated (yellow) or treated with free IMI (0.01 µg/mL, black), aOMVs with an IMI concentration of 0.01 µg/mL (red), or empty OMVs with the same lipid concentration as the aOMVs (white). Each data point represents the mean (n=3) ± standard deviation.

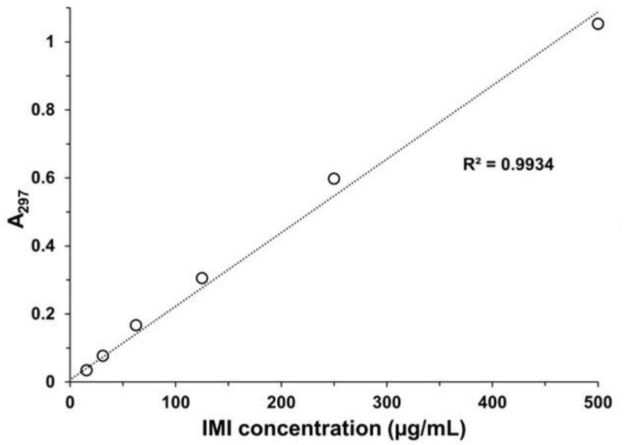


**Figure S6: Characteristic calibration curve for IMI used to determine the concentration of unloaded antibiotic.** The curve was fit using a linear regression, and the coefficient of determination ($R^2$) is displayed.
